# The histone deacetylase *clr3* regulates secondary metabolite production and growth under oxidative stress conditions in *Penicillium brasilianum*

**DOI:** 10.1101/2020.05.01.072108

**Authors:** Daniel Yuri Akiyama, Marina Campos Rocha, Jonas Henrique Costa, Iran Malavazi, Taícia Pacheco Fill

## Abstract

Most of the biosynthetic gene clusters (BGCs) found in filamentous fungi are silent under standard laboratory cultivation conditions due to the lack of expression triggering stimuli, representing a considerable drawback in drug discovery. To access the full biosynthetic potential of these microbes, studies towards the activation of cryptic BGCs are essential. Histone acetylation status is an important regulator of chromatin structure which impacts in cell physiology and, therefore, expression of biosynthetic gene clusters in filamentous fungi. Histone deacetylases (HDACs) and histone acetyl-transferases (HATs) are responsible for maintaining and controlling this process under different cell conditions. In this study, *clr3*, a gene encoding a histone deacetylase in *Penicillium brasilianum* was deleted and associated phenotypic and metabolic changes evaluated. Results indicate reduced growth under oxidative stress conditions in the Δ*clr3* knockout strain. Also, the production of several secondary metabolites including austin-related meroterpenoids, brasiliamides, mycotoxins such as verruculogen and penicillic acid, as well as cyclodepsipeptides was reduced in the Δ*clr3* strain when compared to wild-type strain. Accordingly, addition of epigenetic modulators responsible for HDAC inhibition such as suberoylanilide hydroxamic acid (SAHA) and nicotinamide (NAA) to *P. brasilianum* growth media also culminated in reduction of secondary metabolite production. Mass Spectrometry Imaging (MSI) was applied to compare metabolite production and spatial distribution on the colony. Results suggest that Clr3 plays an important role in secondary metabolite biosynthesis in *P. brasilianum*, thus offering new strategies for regulation of natural product synthesis by assessing chromatin modification in *P. brasilianum*.

## 1. Introduction

Filamentous fungi can produce a vast array of low molecular-weight molecules involved in their secondary metabolism, aiding fungi’s adaptation to environmental conditions, resisting and fighting back predators and competing microbes in their environmental niches (1). These natural products possess a range of bioactive activities, from the treatment of infectious diseases to potent toxic and carcinogenic properties (2). Over the last decades, many efforts have been devoted to identify and study genes involved in the biosynthesis of these compounds due to their important pharmaceutical potentialities (3).

*Penicillium brasilianum* presents a great biosynthetic ability. Metabolites already reported as produced by this species include: diketopiperazines, polyketides, alkaloids, meroterpenoids and cyclodepsipeptides (4). Moreover, this fungus has demonstrated to be an important producer of potently convulsive and bacteriostatic brasiliamides (5), austin-related insecticidal meroterpenes (6) and verruculogen-like tremorgenic alkaloids (7). Functional analysis of *P. brasilianum*’s genome revealed 42 putative biosynthetic gene clusters (BGCs), with 13 clusters related to the biosynthesis of potential polyketide compounds via PKS (polyketide synthase) enzymes; 12 different clusters involved in the production of secondary metabolites formed via NRPS-like enzymes (non-ribosomal peptide synthetases) and 4 hybrid biosynthetic clusters, among them the NRPS-terpene hybrid responsible for alkaloid biosynthesis, indicating the great, yet not fully explored, potential of this organism to produce bioactive secondary metabolites (8).

Bioinformatic, transcriptomic and metabolomic analyses reveal that the majority of microbial BGCs are silent under standard laboratory conditions, due to the absence of triggering stimuli found in nature, representing a huge drawback in drug discovery (9). Thus, novel strategies for the activation of BGCs are essential for natural product prospection.

Multiple factors regulate gene expression, including chromatin packing. Histone modifications play an important role in altering chromatin structure and, therefore, regulating transcription (10). Histone acetylation is the most studied histone modification and depends on the concerted action of histone acetyltransferases (HATs) and histone deacetylases (HDACs) (11). Histone hyperacetylation is known to induce transcriptional activation in several organisms, thus being a solid strategy towards achieving structural diversity of natural products. Enhanced histone acetylation can be achieved by genetic deletion or chemical inhibition of HDACs (12).

Based on the close relation between histone acetylation status and the expression of cryptic BGCs, the objective of this study was to evaluate metabolic profile and phenotypic changes in a *P. brasilianum Δclr3* strain, which is a deletion of the homolog of the class 2 histone deacetylase *hda1* of *Saccharomyces cerevisiae*. Additionally, chemical epigenetic modulation was utilized as an alternative strategy for HDAC inhibition. Secondary metabolism changes were verified through different mass spectrometry-based approaches. Since HDACs regulate BGCs, these results indicate that both genetic manipulation and pharmacological modification of chromatin acetylation are functional approaches to unveil secondary metabolite potential in this *P. brasilianum* allowing further studies in the prospection of novel natural products using this promising fungal model.

## 2. Material and Methods

### 2.1. Fungal strains and culture conditions

The fungus isolation procedure from the root bark of *Melia azedarach* was previously described by Geris dos Santos & Rodrigues-Fo, 2002. *P. brasilianum* (LaBioMMi 136) was cultivated on commercial potato-dextrose-agar (PDA) (Acumedia) and potato-dextrose broth (PD) (Acumedia). Media were autoclaved at 103 KPa (121°C) for 20 minutes. Plates were stored at 30°C for 7 days in darkness. Spores were harvested by washing the agar surface with sterile distilled water and diluted to a final concentration of 10^5^ or 10^6^ spore.mL^−1^.

### 2.2. Genomic DNA extraction

The extraction of fungal genomic DNA was carried out according to the method described by Malavazi e Goldman (13). Briefly, conidia were incubated in 50 mL of commercial potato-dextrose broth (Acumedia) at 25°C, 150 rpm for 72 hours. Mycelia were harvested, ground in liquid nitrogen and suspended in 500 µL of Lysis buffer (200mM Tris-HCl, 250 mM NaCl, 25 mM EDTA, 0,5% [w/v] SDS, pH 8.0). Further genomic DNA purification was performed by phenol/chloroform extraction followed by isopropanol precipitation. Purified DNA was dried and dissolved in ddH_2_O.

### 2.3. Construction of Δclr3 mutant

The *clr3* deletion cassette used in this study was constructed by *in vivo* recombination in *Saccharomyces cerevisiae*, as reported by Malavazi and Goldman (13), 2012. For the cassette construction, fragments of the 5’ and 3’ UTR regions that flank the *clr3* gene were amplified via PCR from the genomic DNA of the wild-type strain (Fig 1). The primers sequences used in this study are listed in S1 Table. Flanking regions contained a small sequence homologous to cloning sites of the pRS426 plasmid. The *hph* gene, which confer resistance to hygromycin, was PCR-amplified from pAN7-1 plasmid and used as a selection marker in the deletion cassette. The three independent fragments, along with *Bam*HI-*Eco*RI-cut pRS426, were transformed into the *S. cerevisiae* FGSC 9721 strain as described by Malavazi and Goldman (13). The plasmids containing the *clr3* deletion cassette were isolated (QIAprep Spin Miniprep Kit) and used as templates to amplify the cassette by using the outermost primers (5F and 3R). All PCR amplifications were performed using Phusion Flash High-Fidelity DNA Polymerase (Thermo Scientific). The *clr3* deletion cassette was transformed into the *P. brasilianum* wild-type strain LaBioMMi 136 (S2 Table) of, according to the protocol described by Malavazi and Goldman (13). Transformants were analyzed by diagnostic PCR and Southern Blot to confirm the insertion at the correct locus.

**Fig 1.**
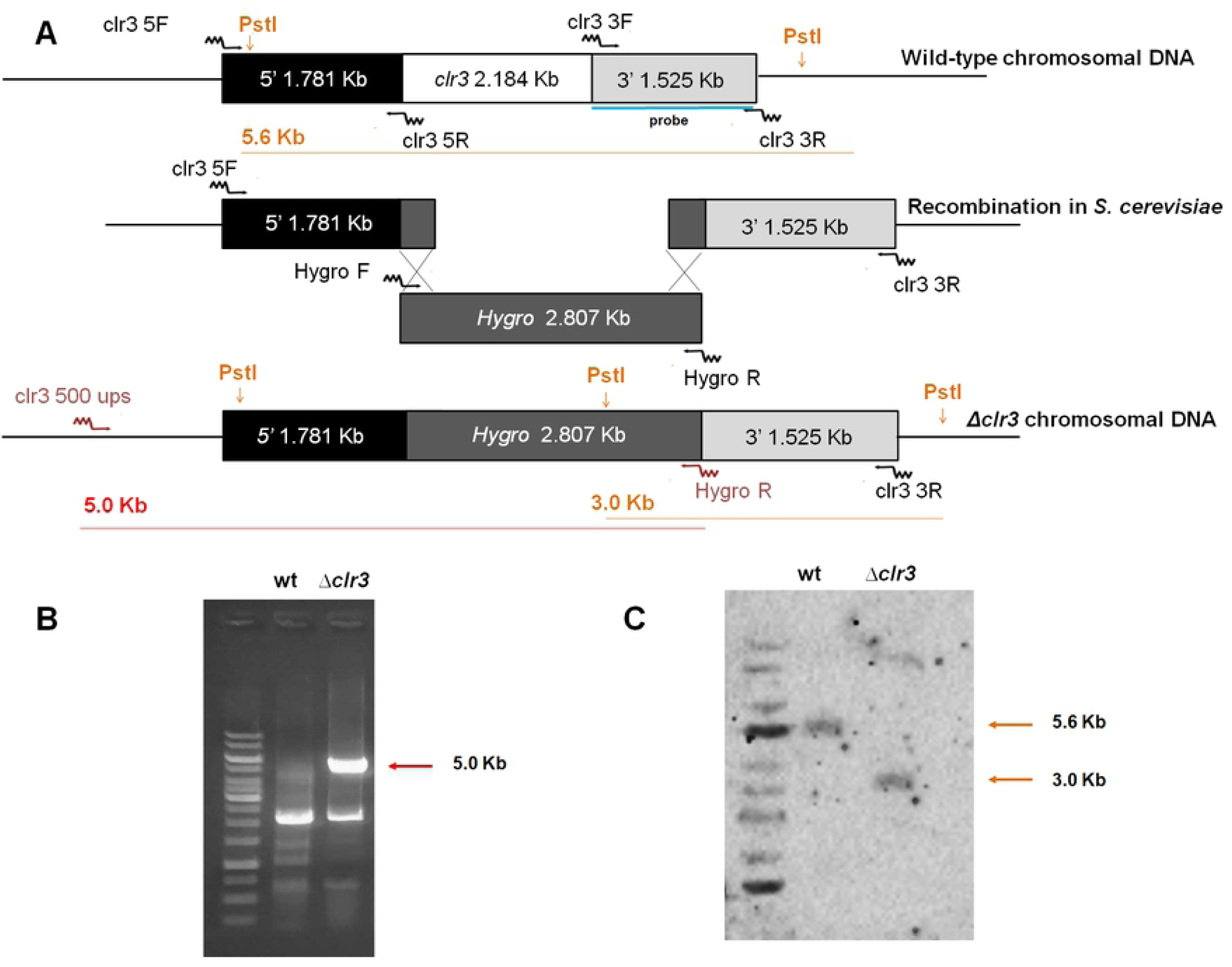
Construction of *clr3* deletion mutant. Gene replacement strategy for *Δclr3* deletion, in which *hph* gene was used as a selection marker. The primer names and annealing regions are indicated by arrows (primer sequences are described in S1Table). (A) Deletion cassettes were constructed by *in vivo* recombination in *S. cerevisiae*. (B) Diagnostic PCR was performed to evaluate *clr3* loci after gene replacement using primers located 500 bp upstream of the deletion cassette, shown in red letters and arrows. (C) Southern blot analysis is shown, probe recognized fragment is indicated by yellow letters and lines.

### 2.4. Southern Blot Analysis

Southern blot analysis was used to show that a single copy of the deletion *clr3*-cassette integrated homologously at the targeted *P. brasilianum clr3* locus. Genomic DNA of the parental *P. brasilianum* and Δ*clr3* strains were isolated as above described was *PstI*-restricted. Chromosomal DNA fragments were separated on a 1% agarose gel and blotted onto Hybond N^+^ nylon membranes (GE Healthcare), following standard techniques (Sambrook and Russell, 2001). Probe labeling for detection was performed using AlkPhos Direct Labeling and Detection System (GE Healthcare) according to the manufacturer’s description. A ChemiDoc™ MP imager (Bio-Rad) was used for gel/blot documentation.

### 2.5. Phenotypic assays for oxidative stress sensibility

To monitor the growth of the *Δclr3* and wild-type strains under oxidative stress 1×10^4^ conidia of each strain were grown in 600 µL of potato-dextrose broth (Acumedia) in 96-well plates supplemented with varying concentrations of paraquat, menadione and H_2_O_2_ (14). Plates were incubated for 72 hours at 30 °C and photographed.

### 2.6. Secondary metabolite extraction

Cultivation was performed for both wild-type and *Δclr3* strains on commercial PDA (Acumedia) and stored at 30 °C for 7 days. After incubation, the petri dish content including solid media and the fungal colony were cut in small pieces and transferred into an Erlenmeyer flask. Extraction was performed using a solvent mixture consisting of methanol, ethyl acetate and dichloromethane (1:2:3). Flasks were sonicated for 30 minutes in ultrasonic bath and vacuum filtered. The extraction process was repeated twice. The solvent was removed under reduced pressure and the final extract was stored at - 20 °C.

### 2.7. UPLC-DAD-MS analysis

The chromatographic system was an ACQUITY™ UPLC system (Waters, Milford, MA, USA) equipped with a diode array detection system. Waters Acquity UPLC BEH C18 analytical column (50 mm × 2.1 mm, 1.7μm) was used as the stationary phase. The mobile phase was composed of 0.1% formic acid (A) and acetonitrile (B). Eluent profile (A/B %): 95/5 up to 2/98 within 8 min, maintaining 2/98 for 5 min and down to 95/5 within 1.2 min. Total run time was 18 min for each run and flow rate, 0.2 mL min^-1^. The injection volume was 5 µL. Mass spectrometry detection was carried out on a Xevo TQD mass spectrometer (Waters Corp., Milford, MA, USA) with electrospray ionization (ESI) source. Analyses were performed in positive ion mode with *m/z* range of 100-1000; capillary voltage at 1.54 kV; source temperature at 149 °C. MassLynx v. 4.1 software was used for data acquisition and equipment control.

### 2.8. High-resolution mass spectrometry analysis

Samples were diluted in methanol. High-resolution mass spectrometry analyses (HPLC-HRMS/MS) were performed in a Thermo Scientific *QExactive© Hybrid Quadrupole-Orbitrap* Mass Spectrometer. Analyses were performed in positive mode with *m/z* range of 115-1500; capillary voltage at 3.4 kV; source temperature at 280 °C; S-lens 100V. The stationary phase was a Thermo Scientific column Accucore C18 2.6 µm (2.1 mm × 100 mm × 1.7 µm). The mobile phase was 0.1% formic acid (A) and acetonitrile (B). Eluent profile (A/B %): 95/5 up to 2/98 within 10 min, maintaining 2/98 for 5 min and down to 95/5 within 1.2 min. Total run time was 25 min for each run and flow rate, 0.2 mL min^-1^. Injection volume was 3 µL. MS/MS was performed by collision-induced dissociation (CID) with *m/z* range of 100–800 and collision energy ranged from 10 to 50V. MS and MS/MS data were processed with Xcalibur software (version 3.0.63) developed by Thermo Fisher Scientific.

### 2.9. Chemical epigenetic modulation experiments

Epigenetic modulation experiments were achieved by using suberoylanilide hydroxamic acid (SAHA) and nicotinamide (NAA) treatments both alone and combined using 48-well microplates. In each well, 1 mL of PD medium, 100 μL of spore solution at 10^6^ spores.mL^-1^ and 5 μL of epigenetic inducers were added to obtain 100 and 200 μM of NAA and SAHA, respectively (final concentration). Cultures were incubated in shaking (70 rpm) for 7 days at 30 °C. Extraction was carried out by liquid-liquid partition after transferring the contents of the wells to separation funnels using ethyl acetate (3 × 2 mL). Organic phase was dried under reduced pressure and final extracts were analyzed by HPLC-HRMS/MS.

### 2.10. Mass Spectrometry Imaging (MSI)

MSI analyses were performed directly on the agar surface using a Prosolia DESI source Modelo Omni Spray 2D^®^-3201 coupled to a Thermo Scientific QExactive® Hybrid Quadrupole-Orbitrap Mass Spectrometer. DESI configuration used was the same set by Angolini et al. (15). The methanol flow rate was set at 10.0 mL.min^-1^. MS data was processed with Xcalibur software (version 3.0.63) developed by Thermo Fisher Scientific. IMS data was acquired using a mass resolving power of 70.000 at *m/*z 200. DESI-MSI data was converted into image files using Firefly data conversion software with a bin width of Δ*m/z* ± 0.03 (version 2.1.05) and viewed using BioMap software (version 3.8.0.4) developed by Novartis Institutes for Bio Medical Research. Color scaling was adjusted to a fixed value for the comparison between the samples.

## 3. Results and discussion

### 3.1. Δclr3 Strain Construction and Phenotypic Analysis

*P. brasilianum* is an important producer of bioactive natural products, such as brasiliamides (16-18), austin-related insecticidal meroterpenes (6,19,20), spirohexalines (20) and verruculogen-like alkaloids (6,7). Recently, Fill et al. reported the draft genome sequence of *P. brasilianum*, revealing a final assembly consisting of a genome size of ∼32.9 Mbp (8). AntiSMASH v3.0 analysis indicated the fungus’ genome presents 42 putative biosynthetic gene clusters (BGCs), 12 of those being non-ribosomal peptide synthetases (NRPSs), 13 polyketide synthases (PKS), 3 terpenes, 2 NRPS/PKS hybrids, 1 hybrid terpene/PKS, and 1 NRPS/terpene (8). This data reveals great secondary metabolite production potential as well as the enzymatic capabilities *P. brasilianum*.

To better access *P. brasilianum’s* cryptic natural products, activation of silent BGCs is one of the approaches to dissect the natural products identity of this organism. Modification of chromatin landscape has been a widely used strategy in order to achieve metabolic diversity in fungi (1). Histone deacetylase activity inhibition, either through gene deletion or epigenetic modulation, has presented relevant results in altering fungal metabolism (11,22) and, in some phytopathogenic species such as *Magnapothe oryzaae*, even pathogenicity properties have been altered (22). Similarly, deletion of *hdaA* gene in *P. chrysogenum* resulted in large expression changes of genes related to pigment production and upregulation of a sorbicillinoids BGC (21), indicating that HDAC inhibition is a feasible strategy in the *Penicillium* genera.

Based on the genome annotation in the NCBI database, four HDACs are present in *P. brasilianum’s* genome. To identify each HDAC, their amino acid sequence was compared through Blast and phylogenetic analysis with the amino acid sequences of known HDACs from *S. cerevisiae, A. nidulans* and *P. digitatum*. A phylogenetic tree was constructed using MEGA6 software based on alignment of amino acid sequences (S1 Fig). HDACs from *P. brasilianum* were predicted based on similarities to the other species’ known HDACs. Altogether, the following HDAC genes were identified: *hosB* (PEBR_24088); *sir2* (PEBR_32801); *clr3* (PEBR_10023) and *rpdA/rpd3* (PEBR_38155).

To evaluate the impact of HDAC activity in *P. brasilianum’s* secondary metabolism and phenotype, the deletion of *clr3* was performed. The *clr3* gene was chosen based on its sequence identity similarity with other known HDACs previously deleted in *A. fumigatus, A. nidulans* and *P. chrysogenum*. The knockout strains for these species presented different fungal development and metabolic profiles (23,24,25), indicating that a *Δclr3 P. brasilianum* strain may also exhibit a different secondary metabolism compared to the parental strain.

Gene knockout was achieved through homologous recombination using a deletion cassette constructed *in vivo* in *S. cerevisiae* using the *hph* gene, encoding hygromycin B phosphotransferase, as a selection marker. Gene deletion strategy, as well as diagnostic PCR agarose gel and Southern Blot analysis can be found in Fig 1 and confirmed the single copy integration of the deletion cassette at the *clr3* locus yielding a null mutant, which was further used for phenotypic and secondary metabolites analyses.

After *clr3* knockout confirmation, phenotypic assays were performed to evaluate possible changes in fungal development in the mutant strain. In both *Trichoderma atroviride* and *Aspergillus nidulans*, deletion of histone deacetylases have led to reduced growth under oxidative stress conditions when compared to their respective parental strains (24,25). The mechanisms underlying oxidative stress response is particularly interesting in phytopathogenic fungi, since most host responses to fungal infection are based on the production of reactive oxygen species from the plant and counteracting responses from the fungus (27). Different *P. brasilianum* strains have already been reported as onion (*Allium cepa* L.) pathogens (28), but little is known about this host-pathogen interaction.

Our investigations revealed that oxidative stress-inducing substances such as H_2_O_2_, paraquat and menadione led to a remarkable reduction of growth in Δ*clr3*. High sensitivity to relatively low concentrations of H_2_O_2_ was observed for the knockout strain (Fig 2), indicating that HDAC inhibition may also be related to *P. brasilianum’s* pathogenic capability, although further pathogenicity assays are necessary to confirm this hypothesis. Oxidative stress is a result of an imbalance between pro-oxidant species and the levels of antioxidant defenses, resulting from the generation of Reactive Oxygen Species (ROS). In contrast to the *Aspergillus ssp*, few data are available on the antioxidative defense system of *P. brasilianum*. For instance, in *A. nidulans*, an expression analysis revealed that CatB, an enzyme responsible for the response to oxidative stress, is regulated with the increase in ROS in wild types, but not in *ΔhdaA* strains (23), suggesting that chromatin modification is part of the regulatory mechanism against oxidative stress. As CatB is one of the known enzymes to be responsible for detoxifying hydroperoxides in hyphae, hypothesizing a positive failure in the positive expression of CatB is one of the main reasons for the sensitivity of *ΔhdaA* strains against an ROS (29). However, to determine the relationship of Clr3 in *P. brasilianum* with enzymes related to oxidative stress detoxification more studies are needed.

**Fig 2.**
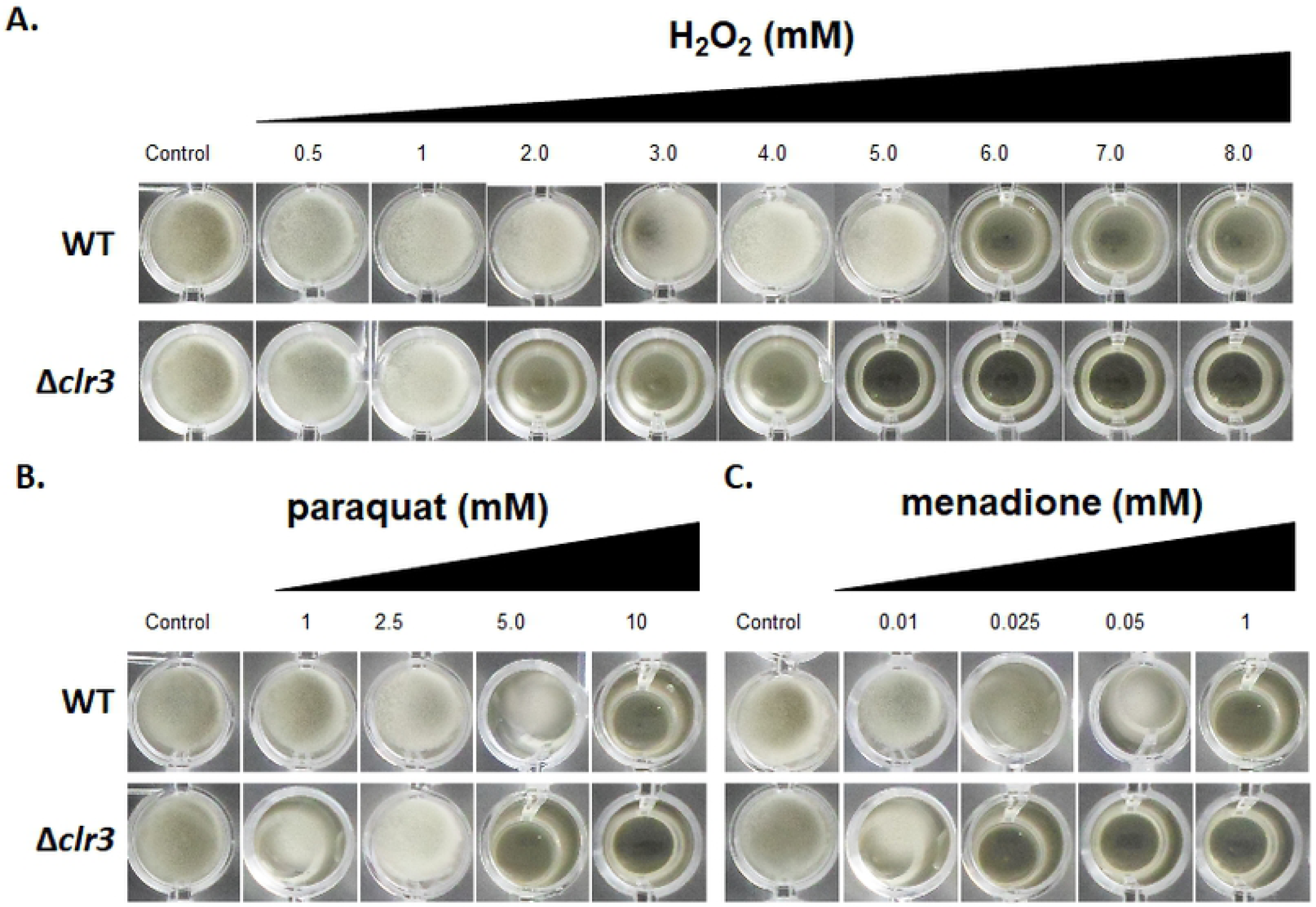
Clr3 null mutants exhibited sensibility to oxidative stress caused by H_2_O_2_, paraquat and menadione. 1×10^4^ conidia of wild-type and mutant strains were inoculated in 600 µL of PD broth (96 well plates) supplemented or not with varying concentration of (A) H_2_O_2_, (B) paraquat and (C) menadione. Plates were incubated at 30 °C for 72h and then photographed.

### 3.2. Natural product diversity in Δclr3 strain

To probe further the contribution of *clr3* in *P. brasilianum* physiology we subsequently used the Δ*clr3* mutant to investigate the secondary metabolite production as an interesting approach to natural product discovery in this species. For metabolic profile comparison, wild-type and *Δclr3* strains were grown in identical cultivation conditions and crude extracts were analyzed by UPLC-MS. Resulting chromatograms (Fig 3) were plotted to the same scale for better comparison.

**Fig 3.**
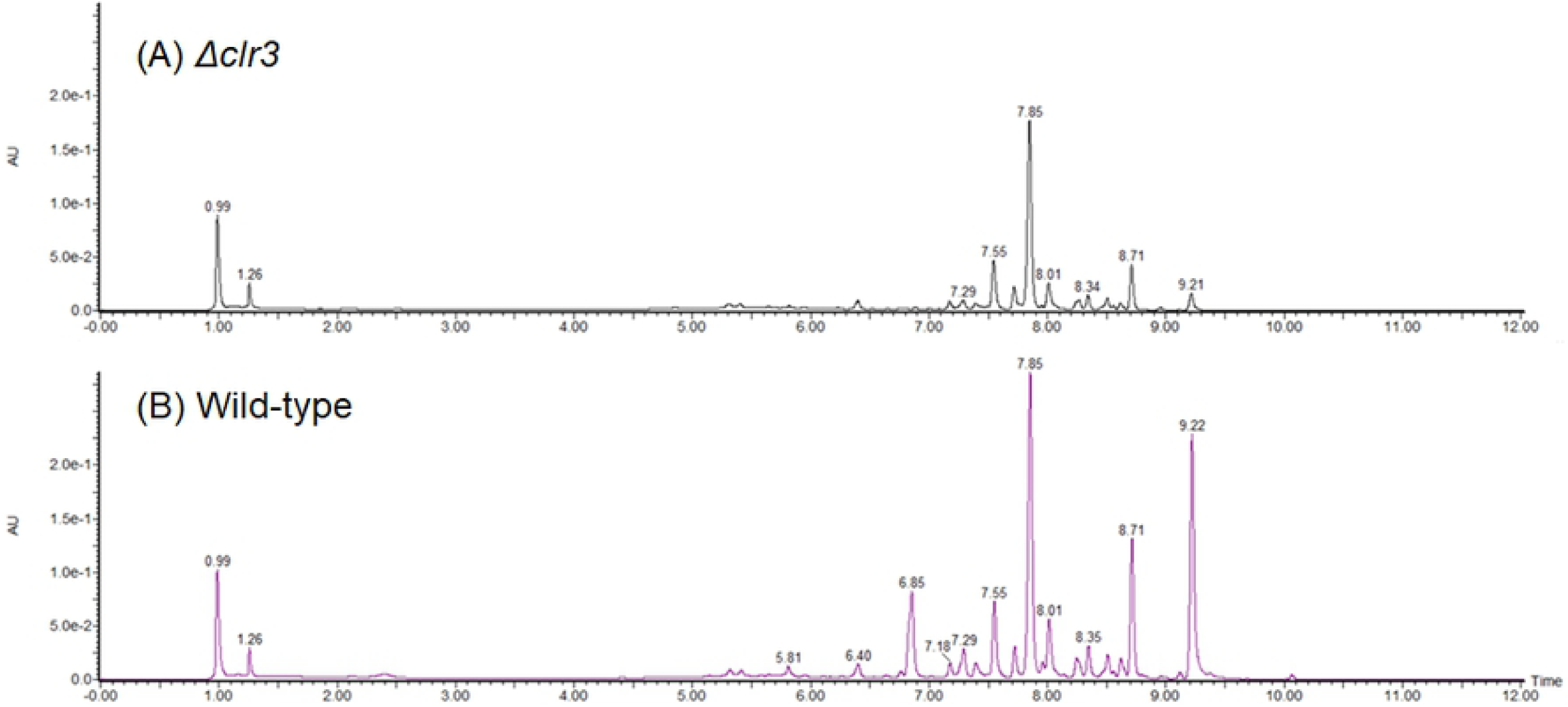
UPLC-DAD chromatograms obtained for the crude extracts from (A) *Δclr3* and (B) wild-type strains of *P. brasilianum*. Chromatograms were plotted to the same scale.

We observed no alteration in the chromatogram profile between the two strains indicating that the deletion of *clr3* did not induce the production of new metabolites or repression of those constitutively formed in our culture conditions. On the other hand, peak areas were significantly different for both strains, indicating that *clr3* has an important regulatory role in secondary metabolite production.

To identify the molecules with different production levels in both strains, crude extracts were further analyzed by HPLC-HRMS/MS and natural product identification was carried out by manually searching on Natural Products databases. Furthermore, the obtained data was compared to HRMS data from previous studies related to *P. brasilianum’s* secondary metabolism (6,7,16-21,30-33). Interestingly, a total of 15 differentially produced compounds already known to be produced by *P. brasilianum* were identified. Data for all metabolites described can be found in Table 1, Fig 4 and S2-S18 Figs.

**Table 1.** HRESI-MS data obtained for compounds 1 – 12.

| Nº | Molecule | Ion formula<br>([M+H]<sup>+</sup>) | Calculated<br>*m/z* ([M+H]<sup>+</sup>) | Experimental<br>*m/z* ([M+H]<sup>+</sup>) | Error<br>(ppm) | Class |
| --- | --- | --- | --- | --- | --- | --- |
| **1** | Isoaustinone | C<sub>25</sub>H<sub>31</sub>O<sub>6</sub> | 427.2115 | 427.2116 | 0.15 | Meroterpenoid |
| **2** | Acetoxydehydroaustin | C<sub>29</sub>H<sub>33</sub>O<sub>11</sub> | 557.2017 | 557.2017 | -0.03 | Meroterpenoid |
| **3** | Austinol | C<sub>25</sub>H<sub>31</sub>O<sub>7</sub> | 443.2064 | 443.2064 | -0.09 | Meroterpenoid |
| **4** | Dehydroaustin | C<sub>25</sub>H<sub>29</sub>O<sub>8</sub> | 499.1963 | 499.1962 | -0.12 | Meroterpenoid |
| **5** | Austinoneol | C<sub>24</sub>H<sub>31</sub>O<sub>6</sub> | 415.2115 | 415.2115 | 0.08 | Meroterpenoid |
| **6** | Brasiliamide A | C<sub>24</sub>H<sub>27</sub>N<sub>2</sub>O<sub>6</sub> | 439.1864 | 439.1865 | -0.05 | Bisphenylpropanoid<br>amides |
| **7** | Brasiliamide B | C<sub>24</sub>H<sub>27</sub>N<sub>2</sub>O<sub>5</sub> | 423.1914 | 423.1914 | -0.09 | Bisphenylpropanoid<br>amides |
| **8** | Brasiliamide C | C<sub>24</sub>H<sub>27</sub>N<sub>2</sub>O<sub>5</sub> | 423.1914 | 423.1914 | 0.06 | Bisphenylpropanoid<br>amides |
| **9** | Brasiliamide D | C<sub>24</sub>H<sub>29</sub>N<sub>2</sub>O<sub>5</sub> | 425.2071 | 425.2070 | -0.16 | Bisphenylpropanoid<br>amides |
| **10** | Brasiliamide E | C<sub>24</sub>H<sub>27</sub>N<sub>2</sub>O<sub>4</sub> | 383.1965 | 383.1966 | 0.24 | Bisphenylpropanoid<br>amides |
| **11** | Verruculogen* | C<sub>27</sub>H<sub>32</sub>N<sub>3</sub>O<sub>6</sub> | 494.2286 | 494.2287 | 0.28 | Diketopiperazines |
| **12** | Verruculogen TR-2 | C<sub>22</sub>H<sub>28</sub>N<sub>3</sub>O<sub>6</sub> | 430.1973 | 430.1969 | -0.84 | Diketopiperazines |
| **13** | Penicillic acid | C<sub>8</sub>H<sub>11</sub>O<sub>5</sub> | 171.0652 | 171.0652 | 0.20 | Polyketide |
| **14** | JBIR 114 | C<sub>30</sub>H<sub>40</sub>N<sub>5</sub>O<sub>7</sub> | 582.2922 | 582.2922 | -0.08 | Cyclodepsipeptides |
| **15** | JBIR 115 | C<sub>30</sub>H<sub>40</sub>N<sub>5</sub>O<sub>7</sub> | 582.2922 | 582.2922 | -0.08 | Cyclodepsipeptides |

\*Verruculogen was identified as a [M+H-H<sub>2</sub>O]<sup>+</sup> adduct in this study

**Fig 4.**
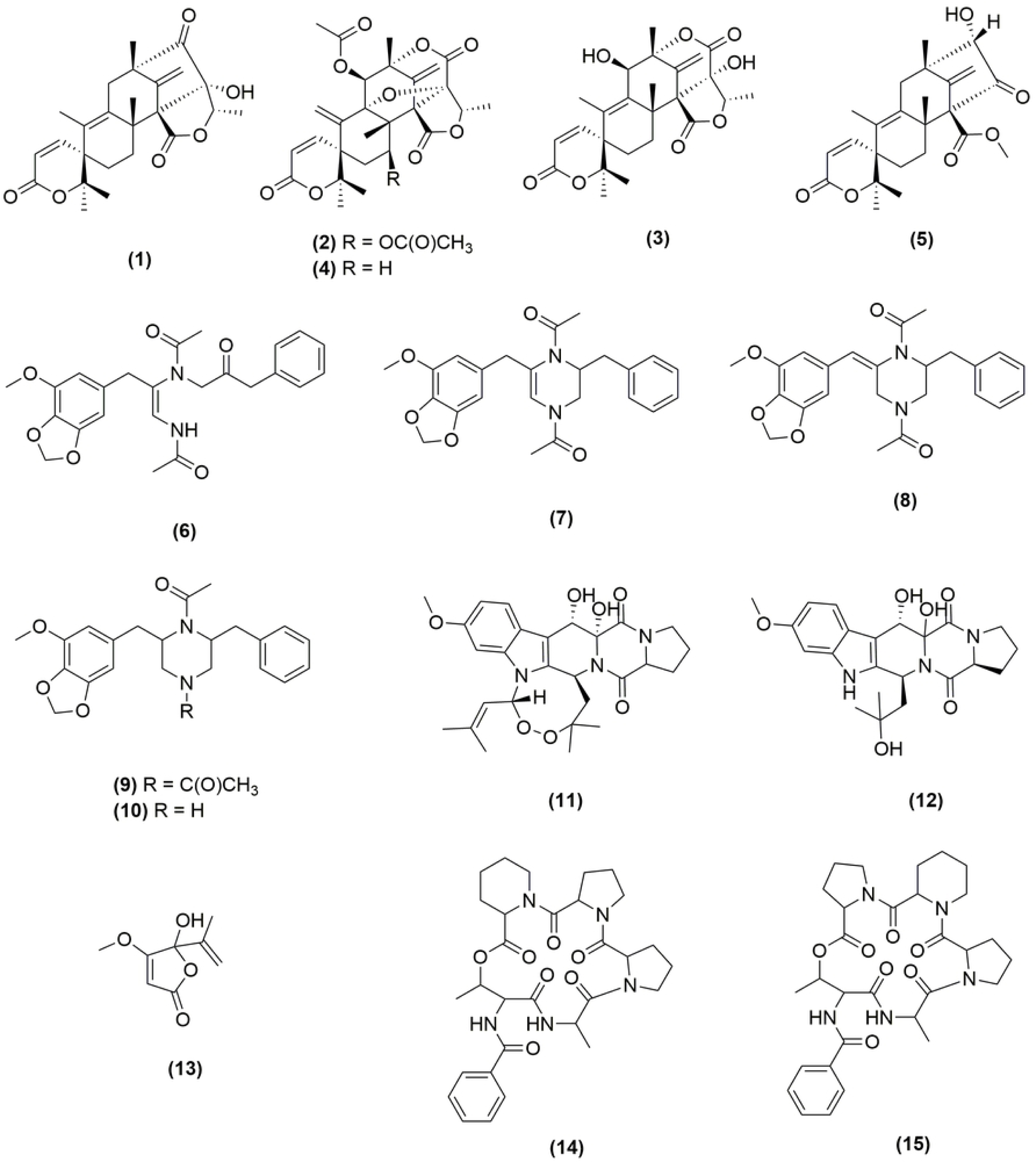
Chemical structures of metabolites identified in this study.

Austin-related meroterpenoids numbered **1**-**5** (Table 1) have been previously related to both *Aspergillus* and *Penicillium* genera, presenting various biological activities such as convulsive and insecticidal (6,18,29,30). Brasiliamides **6-10** are rare examples of fungal phenylpropanoids, whose biosynthesis has not yet been fully elucidated and are produced by *P. brasilianum*, possessing convulsive and a weak antibacterial activity (17,18). Verruculogen **(11)** is a common tremorgenic mycotoxin also produced by both the *Aspergillus* and *Penicillium* genera (6,31). Furthermore, verruculogen derivatives, such as verruculogen TR-2 **(12)**, have also had their production induced in *P. brasilianum* (7). Penicillic acid **(13)** is a polyketide produced by many strains of *Penicillium* fungi, being an important mycotoxin with antibacterial activity (34). Cyclodepsipeptides **14** and **15** present a unique structure with three neighboring cyclic amino acids proline and two pipecolinic acid, indicating a great NRPS-like enzymatic potential in *P. brasilianum* (35).

The lack of novel induced secondary metabolites due to *clr3* deletion was not expected based on its transcription role to suppress gene expression and may be highly related to our cultivation condition, suggesting again the complexity of the secondary metabolite production. Nonetheless, there are several reports in the literature in which perturbation of HDACs activity has led to both up and downregulation of certain BGCs, the suppression of a metabolite or the induction of a new one, indicating a complex response mechanism to chromatin landscape modification in natural product biosynthesis (34,35).

Since the deletion of *clr3* did not induce the production of any new metabolites, different approaches to modulate chromatin structure were sought. Besides gene deletion, HDAC inhibition can be achieved through chemical epigenetic modulation (1). In *Aspergillus niger*, growth in media supplemented by suberoylanilide hydroxamic acid (SAHA), a HDAC inhibitor, was able to alter its secondary metabolism and induce a new pyridone (38). In the *Penicillium* genera, the same strategy was applied in *Penicillium mallochii*, resulting in two new natural sclerotioramine derivatives (39).

In this study, two chemical epigenetic modulators were used: nicotinamide (NAA) and suberoylanilide hydroxamic acid (SAHA) as well as a mixture of both. Extracts from the fungus grown in the presence of these epigenetic modulators were analyzed through HPLC-HRMS/MS. Previously dereplicated compounds **5, 6, 12** and **13** were also identified in these experiments, although production levels varied through the different modulators.

### 3.3. HDACs regulatory role in Penicillium brasilianum secondary metabolism

To better understand the magnitude of HDACs regulation of the secondary metabolism in *P. brasilianum*, as well as to compare HDAC inhibition through *clr3* deletion and chemical epigenetic modulation, metabolite quantification was performed. Metabolites **1-15** were quantified, although data is presented only for those with significant statistical difference (p<0.05). Relative quantification was carried out by comparing peak areas of the identified metabolites in the HPLC-HRMS/MS analyses of the crude extracts from both strains (Fig 5), as well as the ones obtained from the chemical epigenetic modulation experiment (Fig 6).

**Fig 5.**
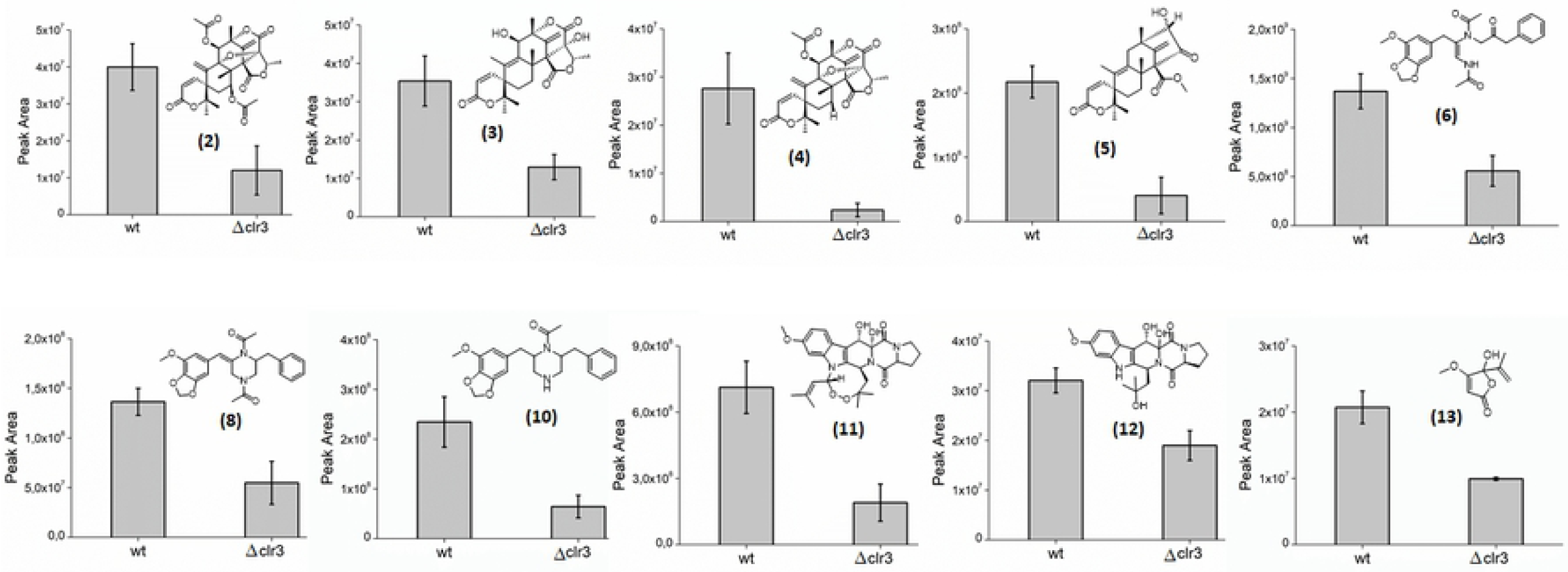
Relative quantification of metabolites 2-8, 10-13 in both wild-type and *Δclr3* strains. The data represent the average value of two replicates. The error bars represent the standard deviation, p ≤ 0.05.

**Fig 6.**
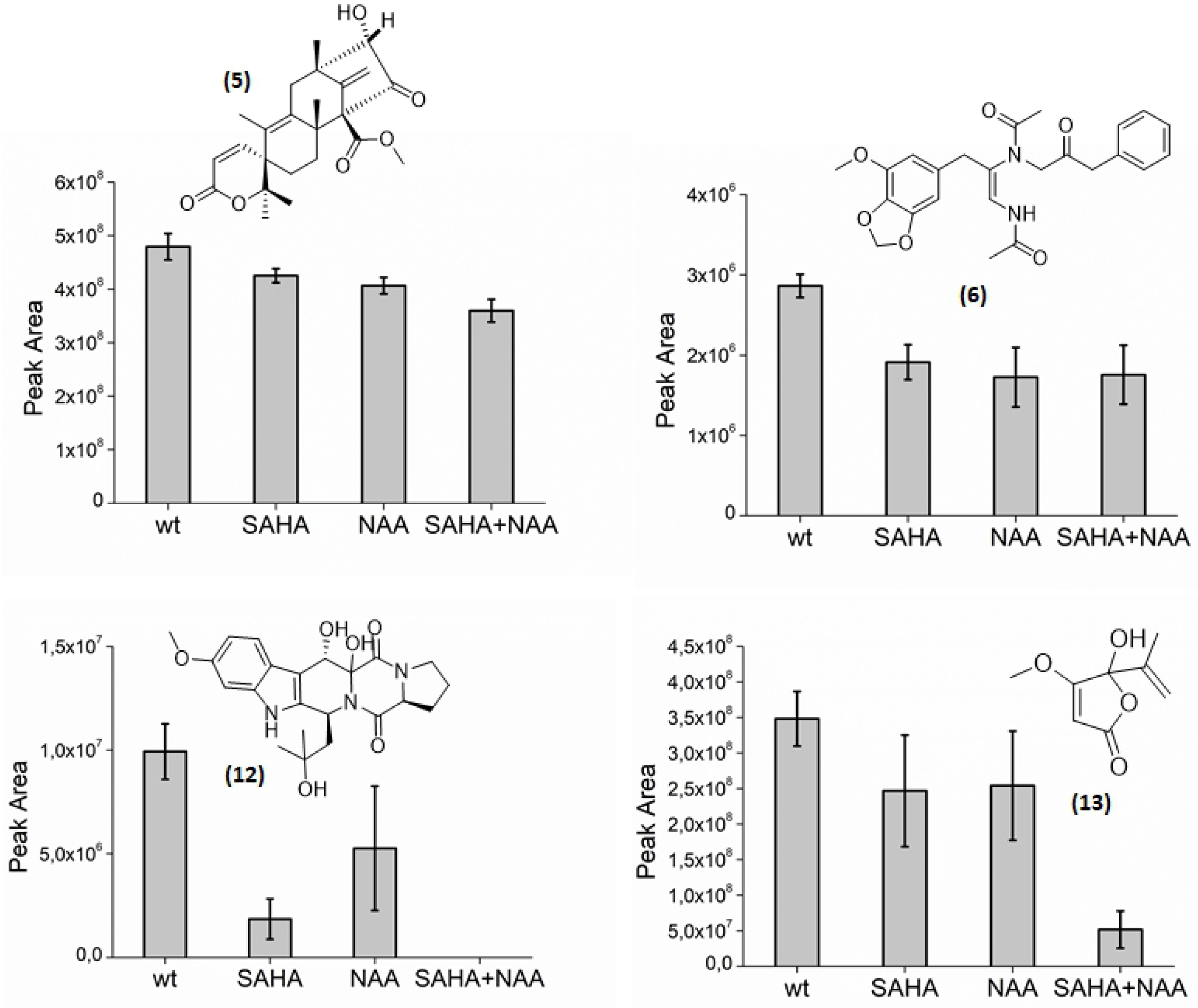
Relative quantification of metabolites 5, 6, 12 and 13 in extracts from *P. brasilianum* grown in the presence of suberoylanilide hydroxamic acid (SAHA), nicotinamide (NAA), as well as a mixture of both (SAHA+NAA). The data represent the average value of two replicates. The error bars represent the standard deviation, p ≤ 0.05.

Comparing the secondary metabolism of both wild-type and *Δclr3*, significant production level differences can be noted. Metabolites **2**-**8, 10, 11** and **13** presented the greatest production suppression, revealing a close relationship between HDAC inhibition and the downregulation of their BGCs. Notably, brasiliamide A **(6)** was the most produced amide in both extracts, however, its biological role has not yet been fully unveiled. This is the first time brasiliamide’s production was evaluated in HDAC inhibition conditions.

Similarly, deletion of SntB, a global histone deacetylase inhibitor in *A. flavus* resulted in the downregulation of the flavotoxin BGC (40). In our study, HDAC inhibition also caused the downregulation of mycotoxins, such as verruculogen **(11)** and penicillic acid **(13)**. On the other hand, metabolites **1, 9, 12, 14** and **15** presented remarkably similar production levels, indicating low correlation between Clr3 activity and the biosynthesis of these natural products.

In *A. fumigatus*, wild-type, *ΔhdaA* mutants and over-expression *ΔhdaA* complement strains exhibited significant differences in secondary metabolism profile, which also resulted in altered virulence properties (24). Ethyl acetate extracts from each strain were added to macrophages and the over-expression strain induced higher cell death at equivalent biomass (24). Albeit the metabolism of *P. brasilianum Δclr3* strain was not evaluated *in vivo*, metabolic changes might alter its endophytic and pathogenic properties as well, although further studies are necessary to evaluate this proposition.

Regarding to the chemical HDAC modulators, the most significant differences were observed in the verruculogen TR-2 concentration **(12)**, in which the mixture of modulators caused a total suppression of its production. Penicillic acid **(13)** and brasiliamide A **(6)** also presented significant differences, especially under the combination of SAHA and NAA.

As an alternative approach to validate these findings, Mass Spectrometry Imaging (MSI) was used to confirm metabolic production differences in **6** and **11**, two most produced metabolites in *P. brasilianum* in this study (Fig 7), as well as to monitor spatial distribution of both molecules in the fungal colony.

**Fig 7.**
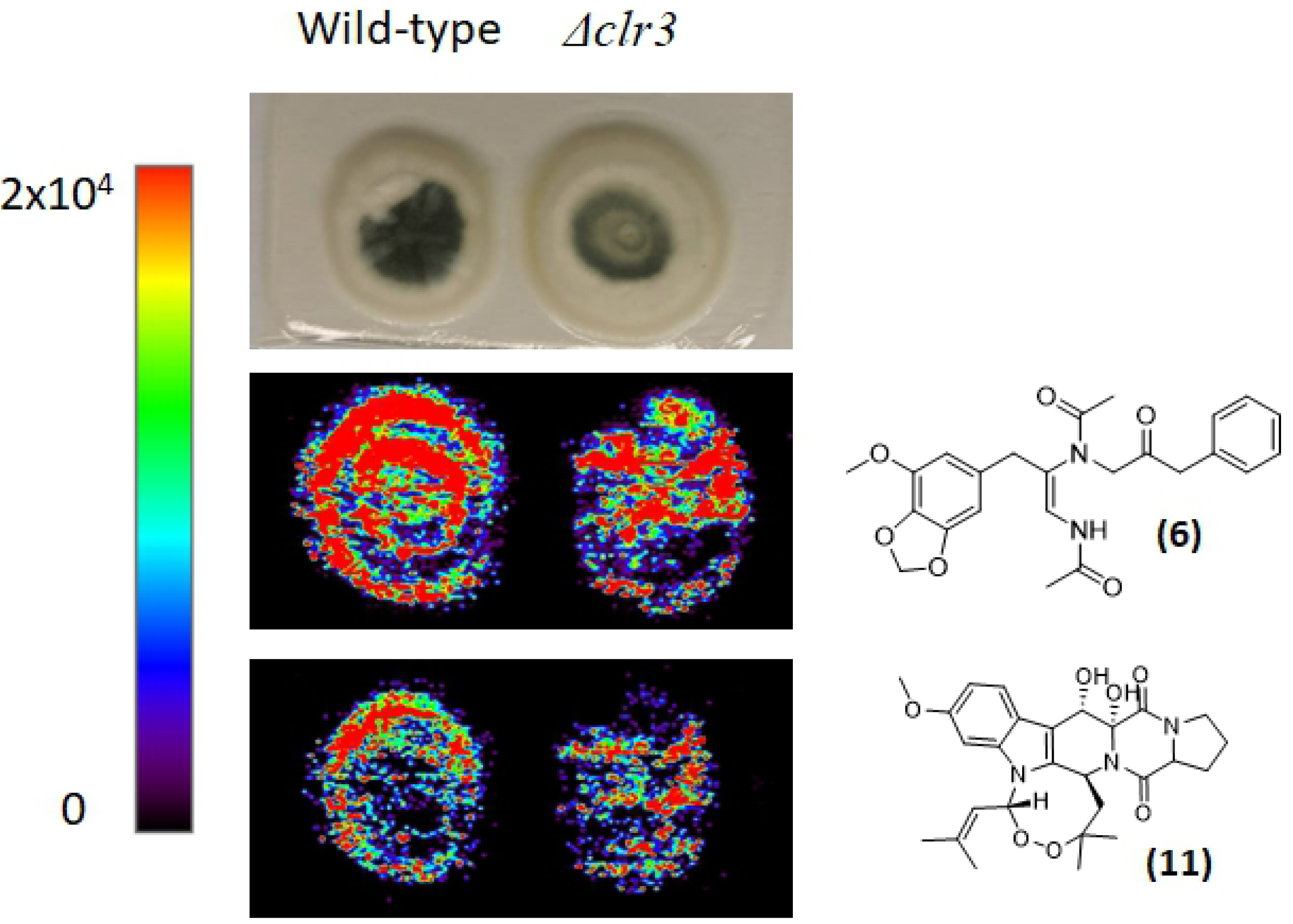
(+) DESI-MSI showing different spatial distributions and concentrations of 6 and 11 on fungal surface. Images are plotted on the same color scale from 0 (black) to 2×10^4^ (red), ion concentration cannot be compared across images due to ionization differences between molecules.

MSI is a powerful tool capable of imaging a vast array of molecules such as metabolites, peptides, lipids and proteins (41), being applied here to provide visualization of spatial distribution of metabolites **6** and **11** (S17 and S18 Figs). DESI-MSI analyses of wild-type and *Δclr3* strains indicated a lower accumulation of **6** and **11** in the colony surface of *Δclr3* than in wild-type. Also, the MSI images indicate that the production of these secondary metabolites occurs in both the conidial and hyphal tissue, since the detection occurred throughout the colony including areas of lower conidiation.

Based on metabolomic approaches, it was possible to verify a close relation between Clr3 activity and secondary metabolite production in *P. brasilianum*. Both the deletion of *clr3*, as well as chemical epigenetic manipulation have led to a downregulation of secondary metabolite production, indicating that histone deacetylases play an important role in regulating the *P. brasilianum* secondary metabolism regulation. To better understand the molecular role of HDACs in the expression of BGCs, further transcriptomics studies are necessary to unveil which BGC are more directly contributing to the phenotypes we described here.

## 4. Conclusions

Understanding filamentous fungi secondary metabolism and its regulation by chromatin structure is an important step towards natural product discovery. Here, we have demonstrated for the first time the effect of HDAC inhibition in *Penicillium brasilianum’s* development and secondary metabolite production. Based on metabolic approaches, both the deletion of *clr3* and epigenetic modulation caused the reduction in production of secondary metabolites such as austin-related meroterpenoids, brasiliamides, verruculogen, penicillic acid and cyclodepsipeptides. In terms of fungal development, *Δclr3* strain exhibited particular sensitivity in growth under oxidative stress conditions. Lastly, this study contributes to a better understanding of HDAC’s role in regulating BGC expression in *P. brasilianum*.

## Acknowledgements

We would like to thank Dr. Edson Rodrigues-Filho for donating the *Penicillium brasilianum* (LaBioMMi 136) strain.

## Supporting information

**S1 Fig. Neighbor-joining phylogenetic tree of HDACs from *S. cerevisiae, A. nidulans, P. digitatum* and *P. brasilianum***. Bootstrap values are indicated on the node of each branch. Gene deleted in this study is marked in red.

**S2 Fig. HRESI-MS data for isoaustinone (1)**.

**S3 Fig. HRESI-MS data for acetoxydehydroaustin (2)**.

**S4 Fig. HRESI-MS data for Austinol (3)**.

**S5 Fig. HRESI-MS data for dehydroaustin (4)**.

**S6 Fig. HRESI-MS data for Austinoneol (5)**.

**S7 Fig. HRESI-MS data for brasiliamide A (6)**.

**S8 Fig. HRESI-MS data for brasiliamide B (7)**.

**S9 Fig. HRESI-MS data for brasiliamide C (8)**.

**S10 Fig. HRESI-MS data for brasiliamide D (9)**.

**S11 Fig. HRESI-MS data for brasiliamide E (10)**.

**S12 Fig. HRESI-MS data for verruculogen (11)**.

**S13 Fig. HRESI-MS data for verruculogen TR-2 (12)**.

**S14 Fig. HRESI-MS data for penicillic acid (13)**.

**S15 Fig. HRESI-MS data for JBIR 114 (14)**.

**S16 Fig. HRESI-MS data for JBIR 115 (15)**.

**S17 Fig. Mass spectrum of ion [M+H]+ *m/z* 439.1878 obtained for brasiliamide A (6) through DESI-IMS**.

**S18 Fig. Mass spectrum of ion [M+H]+ *m/z* 494.2287 obtained for verruculogen (11) through DESI-IMS**.

**S1 Table. Primers used in this study for construction of Δ*hdaA* strain**.

**S2 Table. *Penicillium brasilianum* strains used in this study**.

